# RAD54L promotes nascent DNA degradation and radial chromosome formation in FANC-deficient cells

**DOI:** 10.64898/2026.05.13.724916

**Authors:** Zane Tolbert, Samantha Reed, Steven Goodson, Jennifer M. Mason

## Abstract

Interstrand crosslinks are cytotoxic lesions that inhibit essential processes including replication and transcription. Replication fork reversal occurs in response to interstrand crosslink inducing drug, MMC, but how replication fork reversal promotes repair of interstrand crosslinks is poorly understood. Here, we investigated the role of the RAD54L translocase in interstrand crosslink repair. We found RAD54L is required to promote nascent DNA degradation in FANCD2 and FANCA-depleted cells consistent with a previous study indicating RAD54L promotes replication fork reversal. We further show RAD54L activity is required for formation of radial chromosomes in FANCD2-deficient cells suggesting fork reversal may be required to generate the intermediate undergoing aberrant fusion in FANC-deficient cells. Finally, we demonstrate FANCD2 foci accumulate and DSBs persist in RAD54L-deficient cells indicating RAD54L is required for efficient repair of DSBs. Together, our results indicate RAD54L plays multiple roles in efficient processing and repair of interstrand crosslinks.

## INTRODUCTION

DNA interstrand crosslinks (ICLs) are covalent linkages between bases on opposing strands and inhibit essential processes such as replication and transcription. ICLs can form as a byproduct of metabolic processes including lipid peroxidation as well as exposure to chemotherapeutic drugs such as mitomycin C (MMC).^1^ ICL repair involves the coordination of several DNA repair pathways including nucleotide excision repair, Translesion synthesis (TLS), and homologous recombination.

Central to the repair of ICLs is the Fanconi anemia (FA) pathway comprised of at least 23 proteins that are involved in multiple steps of ICL repair. FANC proteins can be subdivided based on their role in ICL recognition and repair.^2^ The FA core complex is a large E3 ubiquitin complex consisting of FANC proteins including FANCA and FANCC as well as interacting proteins FAAP20 and FAAP100. The primary function of the FA core complex is to recruit FANCT (UBE2T), an E2-ubiquitin conjugating enzyme that monoubiquitinates two FANC proteins, FANCD2 and FANCI, that form a heterodimer (ID complex).^3–6^ The ID complex localizes to DNA damage sites and coordinates downstream processing and repair of the ICL.^7,8^ First, nucleases including ERCC1-XPF (FANCQ) and SNM1A cut one strand on both sides of the ICL leading to unhooking.^8–11^ Translesion DNA synthesis bypasses the unhooked lesion. Finally, homologous recombination factors including BRCA2 (FANCD1), RAD51 (FANCR), and BRCA1 (FANCS) repair the resulting DSB resulting from ICL incision.^2^ In addition, FA proteins including BRCA2, FANCD2 and FANCA proteins protect stalled replication forks from degradation by nucleases including MRE11.^12,13^

Mutations in any of the 23 FA genes results in genome instability, bone marrow failure, growth development and cancer predisposition.^1^ Due the critical role of FA proteins in ICL repair, cells derived from FA patients are hyper-sensitive to interstrand crosslinking drugs including mitomycin C (MMC) and diepoxybutane (DEB). One of the hallmarks of FA is a significant increase in ICL-induced radial chromosomes, aberrant chromosomes resulting from fusion of non-homologous chromosomes by Polθ-mediated end joining.^14^

In response to genotoxic stresses including treatment with MMC, fork reversal occurs whereby the newly synthesized strands anneal to convert a 3-way replication fork into a 4-way junction representing a Holliday junction.^15^ There are two genetically distinct fork reversal pathways. In one pathway, fork reversal is mediated by HLTF, SMARCAL1, and ZRANB3, while the other pathway is mediated by FBH1.^16^ Nascent DNA within the reversed fork is protected from degradation by MRE11, EXO1, and DNA2 by proteins including FANCD2, BRCA2, FANCA, and RAD51.^12,13,17,18^ FANCD2 and BRCA2 only protect forks in the HTLF-dependent pathway, whereas FANCA protects forks in the FBH1-dependent pathway.^16^ Despite being an active area of study, how replication fork reversal promotes ICL repair in human cells is poorly understood.

RAD54L is a member of the SWI2/SNF2 family of proteins that use ATP hydrolysis to translocate on dsDNA, but not ssDNA.^19^ RAD54L works in concert with RAD51 to promote homology directed repair of double strand breaks (DSBs). RAD54L uses ATPase activity to promote RAD51 strand exchange, disassociate RAD51 from heteroduplex DNA after strand exchange has ocurred, and remove RAD51 from undamaged DNA.^20–31^ In mouse and human, RAD54L-deficency results in increased sensitivity to MMC.^32,33^ Recently, RAD54L was shown to promote fork reversal in both HLTF- and FBH1-dependent pathways at replication forks stalled by hydroxyurea (HU).^34^ However, whether RAD54L mediates fork reversal in response to interstrand crosslinks is unknown.

In this study, we investigated the role of RAD54L in ICL repair in response to MMC. We demonstrate RAD54L is required for nascent DNA degradation in FANCD2- and FANCA-depleted cells after treatment with MMC consistent with a role of RAD54L in promoting fork reversal in both HLTF- and FBH1-dependent pathways. We also demonstrate RAD54L knockout rescues MMC-induced radial chromosome formation in FANCD2-depleted cells suggesting RAD54L-mediated fork reversal may be required to generate the ICL repair intermediate that undergoes Polθ-mediated end joining. However, RAD54L-deficiency did not alter hypersensitivity of FANC-deficient cells to MMC. We demonstrate that MMC-induced FANCD2 foci are elevated in RAD54L KO cells and DSBs persist after removal of MMC. These results suggest RAD54L is required for proper resolution of ICLs. Together our results demonstrate RAD54L is required for nascent strand degradation and radial formation in FANC-deficient cells.

## MATERIAL AND METHODS

### Cell lines and drug treatments

U2OS cells were maintained in DMEM media (Gibco Cat #11965–092) supplemented with 10% FBS (EqualFetal, Atlas Biologicals) at 37°C with 5% CO_2_. Generation and characterization of HLTF KO U2OS cells (kind gift from D. Cortez, Vanderbilt University) was previously described.^16^ GM16633 and GM16634 cells (Coriell) were maintained in DMEM media (Gibco Cat #11965–092) supplemented with 10% FBS (EqualFetal, Atlas Biologicals) at 37°C with 5% CO_2_. Hydroxyurea was resuspended in UltraPure distilled water (Invitrogen Cat# 10977-15). Mitomycin C (Caymen Chemical Cat # 11435) was resuspended in DMSO. Drugs were diluted to the concentrations indicated in figure legends.

### Generation of RAD54L knockout U2OS cell line

Guide RNAs flanking exon four of RAD54L were cloned into gRNA cloning vector (Addgene #41824, kind gift from George Church) using Gibson assembly.^35^ Target sequences for the guide RNA were RAD54L gRNA-1 5’ gatgagctgtttagttccca-3’ and RAD54L gRNA-2 5’-gtccagcggtggctgtgttt-3’. U2OS cells were transfected with RAD54L gRNA-1, RAD54L gRNA-2, hCas9 pcDNA3.3-TOPO (Addgene 41815, kind gift from George Church), and pcDNA3.1 zeocin. Single colonies were isolated and expanded after zeocin selection. Genomic DNA was isolated from cells using QuickExtract (Biosearch Technologies) following manufacturers’ instructions. Clones were screened by PCR for deletion of exon four using primers RAD54L_screen_F4 5’-gggagctgtcagagttagc-3’ and RAD54L_sceen_R4 5’ tgagaggtcgaggtgggag 3’. Clones identified by screening were expanded and screened by western blotting with RAD54L antibodies.

### siRNA Transfections

U2OS cells (1.5 × 10^5^/well) were seeded in 6-well plates, cultured for 24 hours, and transfected using Lipofectamine RNAiMAX (Fisher Scientific, 13778150) following manufacturers’ protocol. For negative control samples, Qiagen All-Stars negative control siRNA was used. RAD54L siRNAs were a cocktail of RAD54L siRNA 1-3 from Invitrogen (75nM)^24^. siFANCA was from Dharmacon (J-019283-07-0020, 75 nM) and siFANCD2 was from Ambion Life Sciences (75nM). Target sequences for the siRNAs are listed below:

siFANCA 5’ GGGCCAUGCUUUCUGAUUU
siFANCD2 5’ CAGAGUUUGCUUCACUCUCUA
siRAD54L-1 (HSS112272) 5’ GGAAACCUUUGAGUCAGCUAACCAA
siRAD54L-2 (HSS189050) 5’ UUGGUUAGCUGACUCAAAGGUUUCC
siRAD54L-3 (HSS189051) 5’ GCCAAGGUUGUAGAACGCUUCAAUA

### Replication fiber assay

Cells (40,000) were plated in a 12-well culture plate and cultured for 24 hours. Cells were pulsed with 20µM CldU and 75µM IdU for 20-30 minutes followed by treatment with either 2mM hydroxyurea for four hours or 300nM MMC for one hour. Cells were washed with 1x PBS and collected by trypsinization. Cells were spotted on slides, lysed, and spread as previously described.^36^ Slides were denatured with 2.5 M HCL for 80 minutes. Slides were blocked with 3% BSA in PBS for two hours and incubated with primary antibodies overnight at 4°C to detect IdU (mouse anti-BrdU (BD, 347580, 1:50)) and CldU (rat anti-BrdU (Abcam, ab6326, 1:200)). Slides were washed with 1X PBS and incubated with secondary antibodies chicken anti-rat 488 and rabbit anti-mouse 594 (Invitrogen Alexa Fluor-conjugated antibodies, 1:1000) for one hour at room temperature. Slides were washed with PBS and incubated with tertiary antibodies goat anti-chicken 488 and goat anti-rabbit 594 (Invitrogen Alexa Fluor-conjugated antibodies, 1:1000) for 1 hour at room temperature. Slides were washed with PBS, dried and mounted with Vectashield (VectorLabs). Images were acquired at 60x using a Zeiss Imager M2 epifluorescence microscope with an Axiocam 503 mono camera. For each experiment, CldU and IdU tract lengths were measured for at least 150 replication tracts using ImageJ.

### Immunofluorescence

Cells (4×10^4^) were seeded in 12-well plates containing circular coverslips and treated with 1 μM MMC for 1 hour. Cells were washed with 1X PBS and incubated with HEPES/Triton-X 100 buffer (20 mM HEPES pH 7.4, 3 mM MgCl₂, 50 mM NaCl, 0.5% Triton X-100) for followed by fixation with 3% PFA/3.4% sucrose for 10 min. To detect FANCD2 focus formation in S phase, cells were pulsed with 10 uM EdU for 20 minutes. EdU was detected using Click-It chemistry as per manufacturers’ instructions (Invitrogen). Coverslips were blocked with 3% BSA for 45 minutes and incubated with FANCD2 primary antibodies (Novus Biologicals, NB100-182 1:1000) overnight at 4°C. Cells were washed three times with 1x PBS and incubated with Alexa Fluor-conjugated secondary antibodies for 1 hour at room temp. Slides are then washed three times with 1x PBS followed by dehydration with 70%, 95%, and 100% ethanol. Coverslips were mounted on slides with VectaShield containing DAPI (VectorLabs). At least 50 EdU-positive cells per condition were acquired at 60X magnification using Zeiss Imager M2 epifluorescence microscope with an Axiocam 503 mono camera. the number of FANCD2 foci in each Edu-positive nuclei was counted using Image J.

### Western Blotting

Whole cell extracts were prepared by lysing cells (10^6^/100 ml) in Laemmli buffer (62.5mM Tris-HCl, pH 6.8, 2% SDS, 10% glycerol, 5% 2-mercaptoethanol, 0.002% bromophenol blue) and boiling for 10 min. Extracts were resolved by SDS-PAGE and transferred to PVDF membrane. Membranes are incubated with primary antibodies diluted in 5% milk in in TBST (1X TBS, 0.1% Tween-20) overnight at 4°C. Membranes were washed with TBST three times for 10 min each. Membranes were incubated with HRP-conjugated secondary antibodies (LI-COR, 926-80011/10, 1:2000) for 1 hour at room temperature followed by three washes with TBST for 10 minutes each. Membranes were incubated with WesternSure ECL reagents (Li-COR) and imaged using LI-COR C-DiGit imager with Image Studio software 5.2.5. Primary antibodies used in this study are RAD54L (Santa Cruz Biotechnology, sc-374598 1:500), FANCD2 (Novus Biologicals, NB100-182 1:1000), FANCA (Cell Signaling Technology, 14657S 1:1000), Vinculin (Cell Signaling Technology, 13901S 1:1000), and Tubulin (Novus Biologicals, NB 100-690 1:1000).

### Metaphase chromosome spreads

Cells transfected with the indicated siRNAs were plated in 10 cM dishes and allowed to adhere overnight before adding Mitomycin C. U2OS cells were treated with 150 nM MMC for 24 hours and patient-derived fibroblasts were treated with 300 nM MMC for 24 hours. Cells were washed with DMEM and 1X PBS. U20S cells were treated with 200 ng/ml nocodazole for 16 hours and patient-derived fibroblasts were treated with 300 ng/ml nocodazole for 4 hours. Cells were collected using trypsinization and pelleted by centrifugation. To prepare metaphase spreads, cells were swollen with pre-warmed 0.56% KCl and incubated at 37°C for 30 min. Cells were pelleted by centrifugation at 1000xg for five minutes. Cells are fixed by adding 10mL of ice-cold methanol: acetic acid (3:1) dropwise while gently mixing the tube. Cells were pelted by centrifugation at 1200xg for eight minutes. Fixation with methanol: acetic acid was repeated two times. Fixed cells were dropped on slides to obtain metaphase spreads. Slides were stained with Hoescht 33342 (Thermo Cat #62249.) and acquired at 100x using a Zeiss epifluorescence microscope. At least 50 metaphases for each condition were imaged for each experiment. Metaphases were analyzed and the number of radial chromosomes scored using ImageJ.

### Survival Assays

Cells were seeded at concentrations of 2000, 1000, and 500 cells per well in a 96-well plate. U2OS and GM16633 and GM16634 cells were treated with varying concentrations of MMC for 4 days. Plates were stained with Hoechst 33342 and 1X PBS was added to the wells. Cells were imaged using Molecular Devices ImageExpress Micro 4 and counted using MetaXpress software version 6.7.290.

### Neutral COMET assay

U2OS cells (1.5 × 10^5^/well) were seeded in 6-well plates, cultured for 24 hours, followed by treatment with 1µM MMC for 4 hours. Cells were rinsed with 1X PBS followed by addition of cell culture media. Cells were allowed from for 24 or 48 hours. Cells were collected by trypsination, pelleted and resuspended in 500 µl 1XPBS. Neutral COMET assay was conducted as described.^37^ COMET slides were purchased from R&D Systems (4250-050-K). At least 50 nuclei per condition were acquired at 20x magnification using a Zeiss Imager M2 epifluorescence microscope with an Axiocam 503 mono camera. Tail area was calculated using the open COMET plugin in ImageJ.^38^

### Graphing and Statistical analysis

Results were graphed using GraphPad Prism v.10.4.1. Statistical significance was calculated using the statistical test indicated in figure legends using Prism software.

## RESULTS

### RAD54L promotes replication fork degradation in FANC-deficient cells

A previous study demonstrated RAD54L loss rescued degradation in BRCA2 and 53BP1 depleted cells after treatment with HU due to loss of fork reversal.^34^ FANCD2 and FANCA protect reversed forks by preventing degradation of nascent DNA by nucleases including MRE11.^12,16^ FANCD2 has been shown to protect forks reversed by HLTF and FANCA protects replication forks reversed by FBH1.^16^ To determine how RAD54L impacts fork degradation in FANC-deficient cells, we examined replication fork degradation in RAD54L KO cells with and without depletion of FANCD2 (siFANCD2) or FANCA (siFANCA) (**Fig. 1A, 1B, Sup. Fig. 1A**). Cells were pulsed sequentially with IdU and CldU followed by treatment with MMC. In untreated cells, we observed no significant difference in CldU/IdU ratio in any sample. After MMC treatment, siFANCD2 or siFANCA had a significant reduction in the CldU/IdU ratio in U2OS cells compared to a non-silencing control (siNS) indicating increased nuclease-mediated degradation of nascent DNA. In RAD54L KO cells, we did not observe a significant difference in the CldU/IdU ratio indicating loss of RAD54L does not impact replication fork stability. In contrast to U2OS, siFANCD2 and siFANCA had no impact on the CldU/IdU ratio in RAD54L KO cells indicating that RAD54L is required for fork degradation in FANC-depleted cells. The impact of RAD54L KO on fork degradation in FANC-deficient backgrounds was not specific to MMC as RAD54L KO also rescued fork degradation in FANCA and FANCD2-depleted cells after treatment with HU (**Sup. Fig 2A-C**) indicating RAD54L promotes fork degradation in FANC-deficient cell in response to different genotoxic treatments.

**Figure 1.**
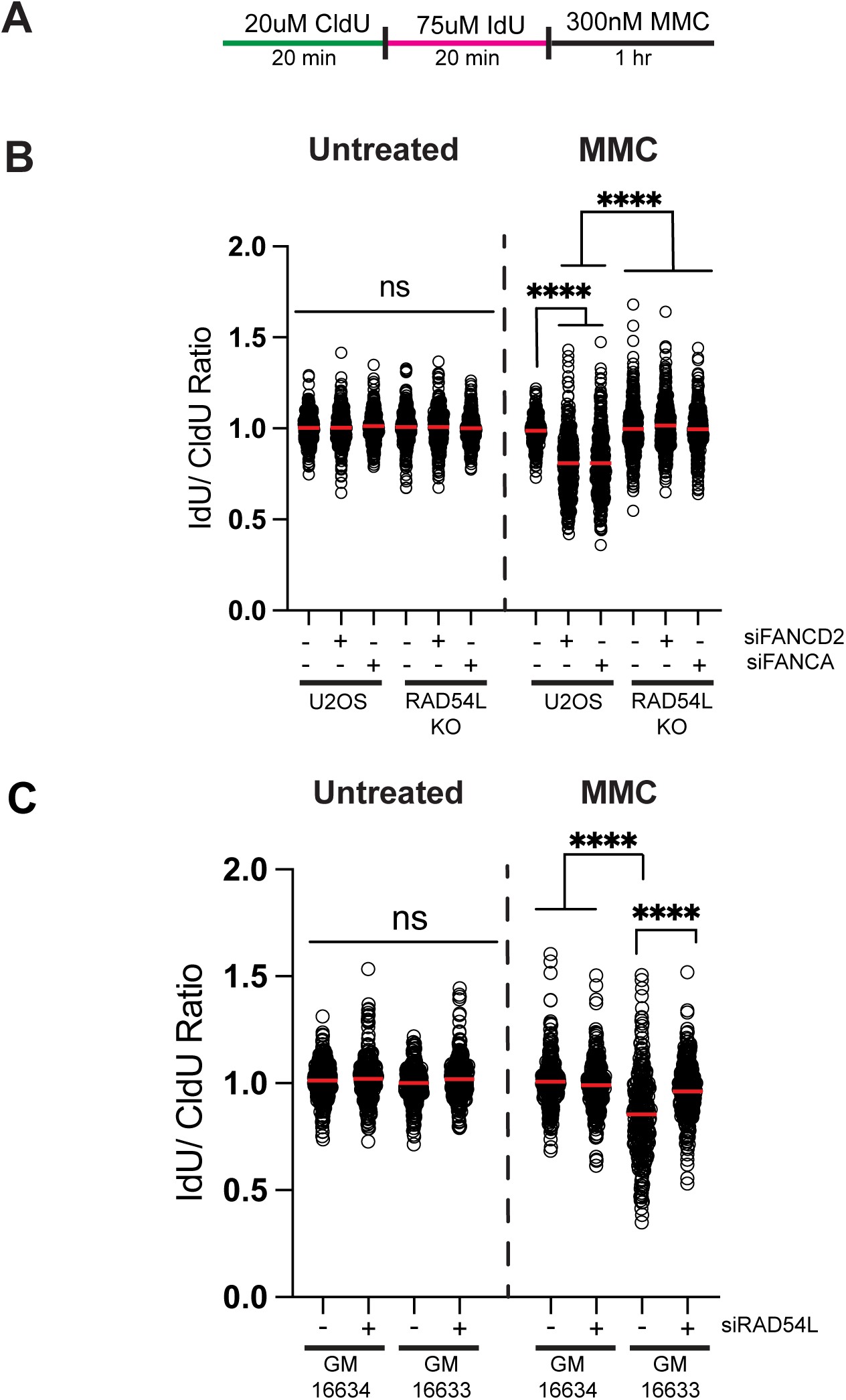
Depletion of RAD54L rescues fork degradation phenotypes in FANCD2 and FANCA deficient cells following treatment with MMC. (A) Schematic of the replication fiber assay. (B) Dot blot showing IdU/CldU ratios of replication tracts. Red line represents the mean. Dot blots depict data from two independent experiments. (C) Dot blot showing IdU/CldU ratios of replication tracts. Red line represents the mean. Dot blots depict data from two independent experiments. \*\*\*\**p*<0.0001, ANOVA, Tukey HSD.

**Figure 2.**
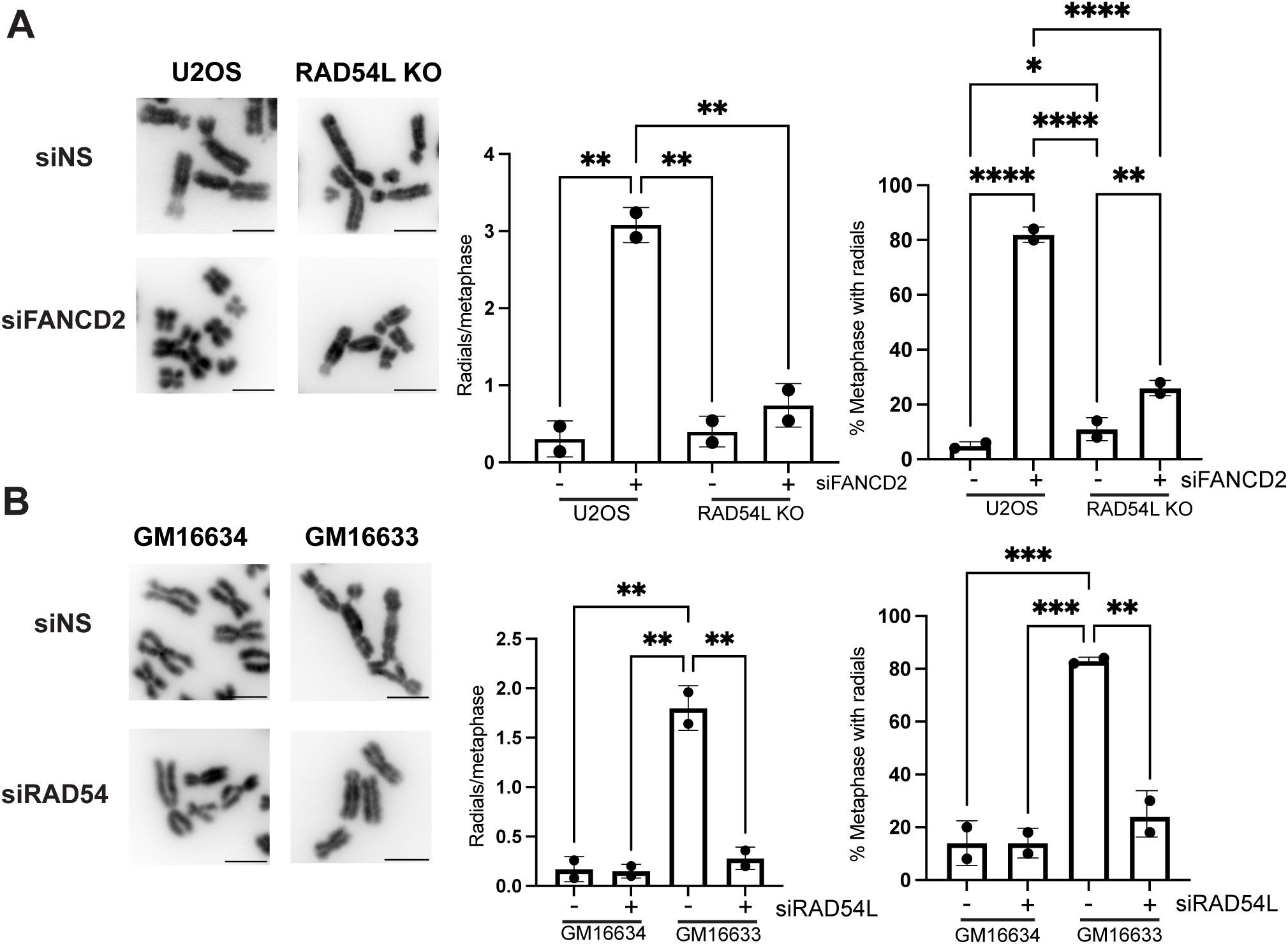
Depletion of RAD54L rescues MMC-induced radial chromosome formation in FANCD2-deficient cells. (A) Representative images of chromosomes for indicated samples treated with MMC. Scale bars equal 5µm. Bar graphs depict average number of radials per metaphase or percentage metaphases with chromosome radial formation. Error bars=STD. N=2. \**p*<0.05, \*\**p*<0.01, \*\*\*\**p*<0.0001, ANOVA, Tukey HSD. (B) Representative images of metaphase chromosomes. Bar graphs depict radials per metaphase and percentage of metaphases with chromosome radial formation. Error bars=STD. N=2 \*\**p*<0.01, \*\*\**p*<0.001, ANOVA, Tukey HSD.

We confirmed RAD54L promotes fork degradation after MMC treatment using FANCD2-deficient fibroblasts derived from a patient with Fanconi Anemia (GM16633) (**Sup. Fig. 1B, Fig 1C**). A FANCD2-complemented cell line (GM16634) was used as a control. We observed no significant difference in the IdU/CldU ratio in untreated samples. FANCD2-deficient cells exhibited a significant reduction in the CldU/IdU ratio compared to FANCD2-complemented fibroblasts indicating increased degradation of stalled replication forks. As observed in U2OS cells, RAD54L depletion (siRAD54L) rescued replication fork degradation in FANCD2-deficient fibroblasts. In addition, RAD54L depletion rescued nascent DNA degradation in FANCD2-deficient fibroblasts treated with HU (**Sup. Fig. 2D**). Together, these results indicate RAD54L promotes nascent strand degradation in FANC-deficient cells after MMC treatment.

### RAD54L promotes MMC-induced radial formation in FANCD2-deficient cells

Next, we determined whether RAD54L was required to promote genome instability in FANC-deficient cells. In response to MMC, FANCD2-deficient cells have an increase in chromosome anomalies including the formation of radial chromosomes.^1^ Thus, we determined the impact of RAD54L KO on MMC-induced radial formation in FANCD2-deficient cells (**Fig. 2A**). In siFANCD2 U2OS cells, metaphases contained on average 3.1 radial chromosomes per metaphase compared to 0.31 in siNS U2OS controls. In addition, 81% of metaphases contained at least one radial. Surprisingly, RAD54L KO rescued MMC-induced radial formation in siFANCD2 cells (0.74 radials/chromosome and 26% metaphases with at least one radial).

We confirmed this result by examining radial formation in FANCD2-deficient (GM16633) and FANCD2-complemented (GM16634) patient fibroblasts (**Fig 2B**). FANCD2-deficient fibroblasts exhibited a significant increase in MMC-induced radials (1.8 radials/metaphase in GM16633 vs. 0.17 radials/metaphase in GM16634) and the percentage of metaphases with at least one radial (83% GM16633 vs 14% GM16634). As observed in U2OS, RAD54L depletion significantly reduced the average number of radials/metaphase (0.28 radials/metaphases in GM16633 siRAD54L vs. 1.8 radials/metaphase in GM16633 siNS) and the percentage of metaphases with at least one radial (83% GM16633 siRAD54L vs. 24% GM16633 siNS). Together, these results indicated RAD54L is required for radial chromosome formation in FANCD2-deficient cells after MMC treatment.

### RAD54L KO cells does not significantly alter hypersensitivity of FANC-deficient cells to MMC

Given the role of RAD54L in promoting radial formation and nascent strand degradation, we examined the impact of RAD54L loss on hypersensitivity of FANC-deficient cells to MMC (**Fig. 3A**). In U2OS cells, RAD54L KO alone did not result in a significant decrease in cell survival after MMC. Previous reports have shown that RAD54L-deficient cells exhibited increased sensitivity to MMC, however, the MMC concentrations used were significantly higher than the concentrations used in these experiments due to the hypersensitivity of FANC-depleted cells to MMC.^33^ Surprisingly, RAD54L KO did not significantly alter hypersensitivity of FANCA- or FANCD2-depleted cells in response to MMC.

**Figure 3.**
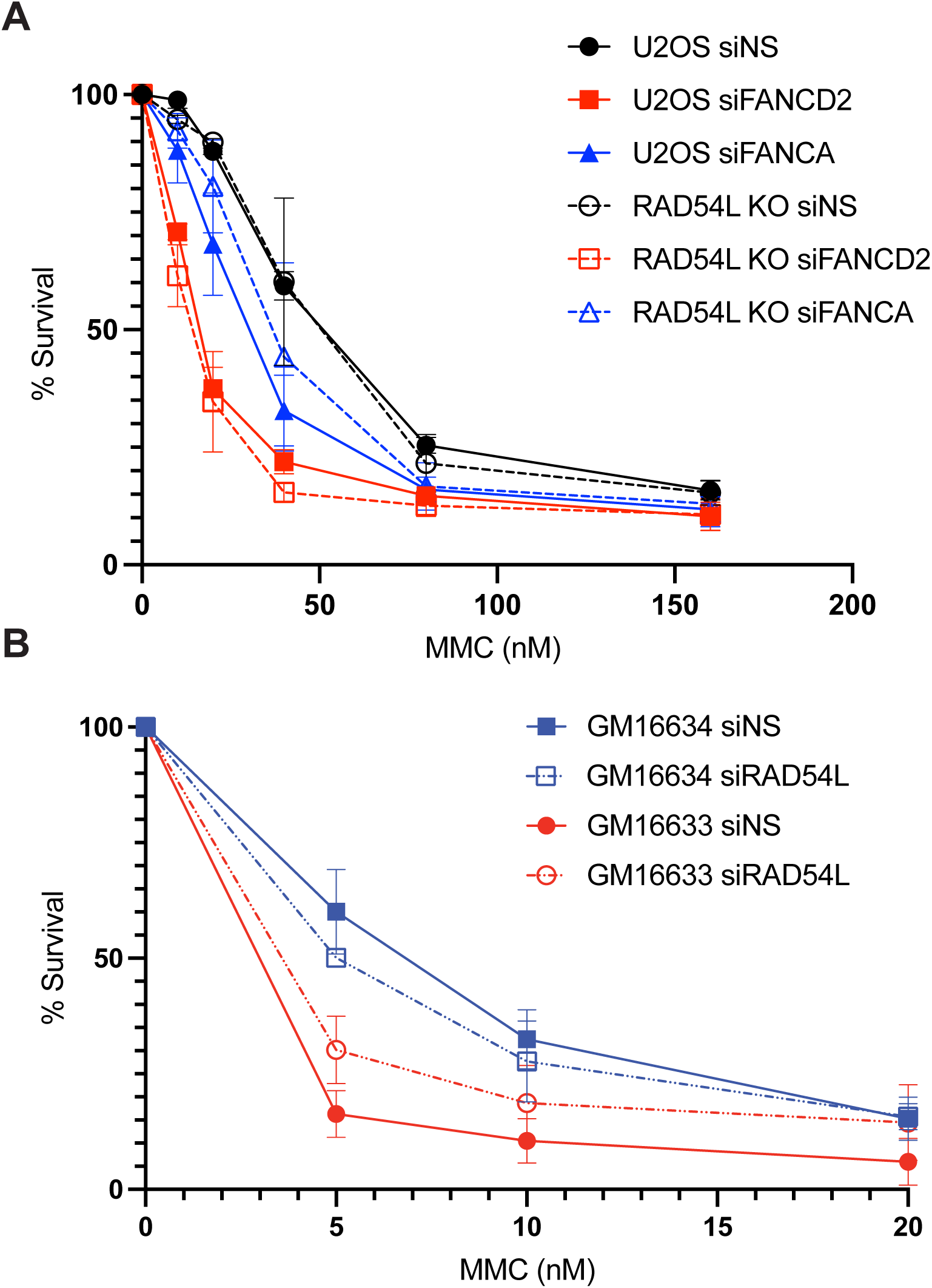
Depletion of RAD54L does not significantly change hypersensitivity of FANC-deficient cells to MMC. (A) Survival curve of U2OS and RAD54L KO cells depleted of FANCD2 or FANCA following treatment with MMC at indicated concentrations. N=2. error bars = SEM. (B) Survival curve of GM16633 and GM16634 cells depleted of RAD54L following treatment with MMC at indicated concentrations. N=2. error bars = SEM.

We obtained a similar result in FANCD2-deficient patient fibroblasts depleted of RAD54L (**Fig 3B).** As expected, FANCD2-deficient cells (GM16633) resulted in hypers-sensitivity to MMC compared to FANCD2-complemented cells (GM16634). RAD54L knockdown in FANCD2-complemented cells resulted in a slight, but insignificant decrease in cell survival. Interestingly, RAD54L depletion led to a slight increase in cell survival in FANCD2-deficent cells suggesting that RAD54L depletion slightly improves survival of FANCD2-deficient cells to MMC. Together, these results indicate RAD54L is not promoting hypersensitivity of FANC-deficient cells.

### FANCD2 foci are increased in RAD54L knockout cells

RAD54L loss rescued nascent strand degradation and radial formation in FANC-deficient cells but did not significantly alter hypersensitivity of FANC-deficient cells to MMC indicating RAD54L may have additional roles in ICL repair. Upon treatment with MMC, FANCD2 is recruited to ICLs and functions as a scaffold to recruit and coordinate downstream repair events^4,8^. We examined induction of FANCD2 foci in control and RAD54L KO cells in response to MMC (**Fig 4**). We pulsed cells with EdU to identify cells in S phase followed by treatment with MMC. After 1 hour of treatment, we observed an increase in FANCD2 foci in U2OS cells compared to untreated controls (**Fig 4A, B**). RAD54L KO cells exhibited a significant increase in FANCD2 focus formation in MMC-treated cells compared to U2OS cells. These results indicate RAD54L is required to limit FANCD2 focus formation suggesting RAD54L KO cells exhibit a defect in ICL repair.

**Figure 4.**
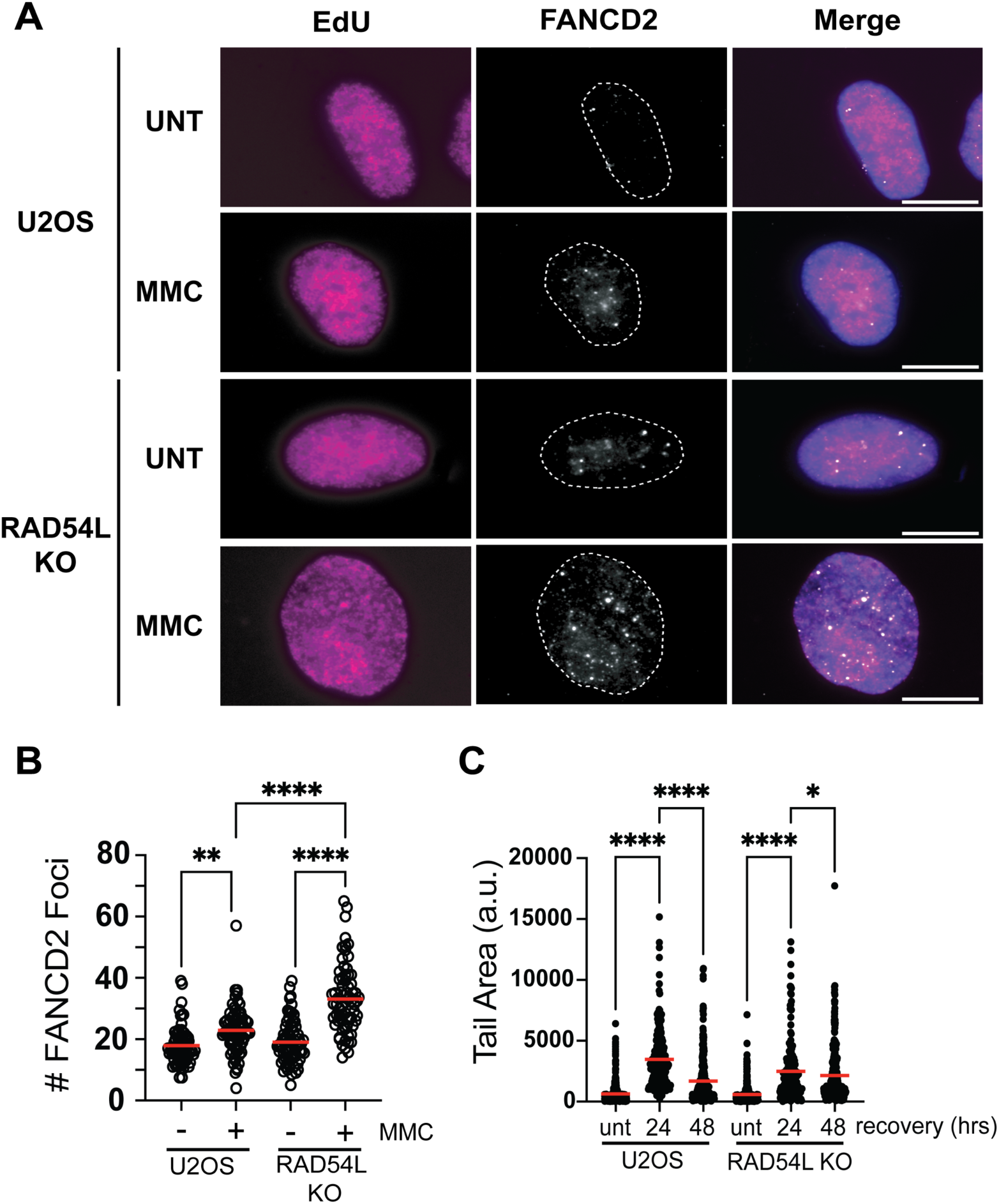
RAD54L is required for efficient resolution of ICLs. (A) Representative images depicting FANCD2 focus formation in the indicated samples after treatment with 1 μM MMC for 1 hour. Scale bar = 10µm. (B) Dot blot represents number of FANCD2 focus formation after the indicated cells. Red line represents the mean. N=3. \*\**p*<0.01,\*\*\*\**p*<0.0001, ANOVA, Tukey HSD (C) Dot blot depicts tail area measurements for the neutral COMET assay after recovery from a 4-hour 1µM MMC treatment. N=3. Red line represents the mean. \**p*<0.05, ****, *p*<0.0001, ANOVA, Tukey HSD.

### Double strand breaks persist in RAD54L KO cells

Persistent and unrepaired ICLs can result in replication forks being cleaved by the MUS81 endonuclease^9^. To determine whether RAD54L is required for resolution of ICLs, we examined DSB formation using the neutral COMET assay (**Fig 4C**). U2OS cells were treated with 1 μM MMC for 4 hours and allowed to recover for 24 and 48 hours. In U2OS cells, DSBs were elevated at 24 hours post MMC treatment, but significantly reduced by 48 hours post MMC indicative of ICL resolution. Although DSBs are induced in RAD54L KO cells 24 hours post MMC treatment, cells exhibit reduced levels of DSBs compared to U2OS cells. Furthermore, DSBs persist in RAD54L KO 48 hours post MMC. Together, these results suggest RAD54L has a role in processing and repair of ICLs.

## DISCUSSION

In this study, we provide evidence that RAD54L promotes both nascent DNA degradation and radial formation in FANC-deficient cells but does not have a significant impact on the hypersensitivity of FANC-deficient cells to ICL inducing agents. Finally, RAD54L limits FANCD2 focus formation after MMC and is required for resolution of DSBs in response to ICLs. Together, these results indicate RAD54L likely has multiple functions during ICL repair.

In response to HU, RAD54L promotes fork reversal and in both the HLTF-and FBH1-dependent pathways.^34^ Here, we have expanded upon these results and demonstrate that RAD54L activity is required to promote degradation of nascent DNA in FANCD2- and FANCA-deficient cells in response to HU and MMC. FANCD2 and FANCA protect nascent DNA from degradation in the HLTF/SMARCAL1/ZRANB3 and FBH1-mediated fork reversal pathways, respectively.^16^ Thus, we propose that RAD54L promotes fork reversal in both pathways in response to MMC indicating RAD54L is promoting fork reversal in response to different genotoxic stresses.

In FANC-deficient cells, radial chromosomes arise due to fusion of non-homologous chromosomes by Polθ-mediated end joining.^14,39^ There are two ICL intermediates that are generated during repair that produce a DNA end that can serve as a substrate for Polθ-mediated end joining. First, reversed forks produce a DNA end that is protected from nuclease-mediated degradation by FANCD2.^12,16^ The deprotected end in FANCD2-deficient cells may serve as the template for Polθ-mediated end joining. Consistent with this model, SCAI depletion leads to fork deprotection and increased Polθ-dependent radial formation.^40^ Given RAD54L is required for FANCD2-mediated fork protection, loss of fork reversal may prevent Polθ-mediated joining of unprotected forks. Second, DSBs are formed during ICL repair after unhooking by ERCC1-XPF and have been proposed to be the intermediate joined by Polθ to generate radial chromosomes.^14^ In Xenopus extracts, replication fork reversal of one of the converged forks is required to generate a substrate suitable for incision by ERCC1-XPF.^41^ Therefore, RAD54L KO may prevent proper ICL processing by nucleases including ERCC1-XPF. Consistent with this, we observed increased FANCD2 focus formation in RAD54L KO cells and a slight, but significant reduction in DSBs suggesting RAD54L is required for proper ICL resolution. Future experiments are warranted to distinguish between these two models.

Our findings that RAD54L KO does not result in significant chromosome anomalies and reduced DSB formation after MMC differs from cellular phenotypes of other fork reversal mutants. ZRANB3 and RAD51-mediated fork reversal is implicated in promoting bypass, or traverse, of ICLs allowing for post-replication repair.^42^ Depletion of ZRANB3 and RAD51 resulted in increased chromosomal anomalies and DSBs due to loss of ICL traverse. There are at least two explanations for the differing results. First, it is possible a subset of fork reversal proteins have different roles in ICL repair. For instance, a subset of fork reversal enzymes such as ZRANB3 may promote ICL traverse, while others such as RAD54L may promote replication-dependent repair by ERCC1-XPF. In Xenopus extracts, fork reversal is required for ICLs repair dependent on ERCC1-XPF, but not for repair of ICLs by NEIL3 glycosylase consistent with fork reversal not having a universal role in all ICL repair pathways.^41^ Second, other proteins may be compensating loss of RAD54L. In mice and humans, RAD51AP1 or RAD54B are able to compensate for RAD54L in response to MMC.^32,33^ Given RAD54L KO rescues nascent strand degradation and radial formation in FANCD2- and FANCA-depleted cells, it is unlikely RAD51AP1 or RAD54B can compensate for loss of RAD54L-mediated fork reversal. Future work will need to determine what functions of RAD54L are required to promote radial formation and ICL repair and whether or not RAD51AP1 or RAD54B can compensate for RAD54L.

Despite rescuing nascent strand degradation and radial chromosome formation in FANC-deficient cells, RAD54L KO had no impact on MMC hypersensitivity of FANC-deficient cells. We propose this is likely due to additional roles of RAD54L in promoting repair of ICLs. In mice, Rad54l is required to repair MMC-induced breaks formed by the nuclease Mus81 and RAD54L-deficient cells exhibit increased sensitivity to higher doses of MMC.^32,33,43^ We observed increased FANCD2 focus formation and persistent DSBs in response to MMC consistent with a role of RAD54L in promoting resolution of ICLs. An interesting future avenue of research is to determine what activities of RAD54L are required for the different functions in ICL repair.

## DATA AVAILABILITY STATEMENT

Request for further information, resources and reagents should be direct to and will be fulfilled by the lead contact, Jennifer Mason

## MATERIAL AVAILABILITY STATEMENT

All research reagents generated during the course of this study will be made available upon request from the lead contact.

## ACKNOWLEDGEMENTS

This research was supported by a grant from the National Institutes of Health (NIH) R35GM142512 to J.M.M

## AUTHOR CONTRIBUTION STATEMENT

Conceptualization, J.M.M.; Investigation, Z.T., S.R., S.G.; Supervision, J.M.M; Writing-Original Draft, Z.T., S.R, J.M.M.; Writing-review and editing, Z.T., S.R., J.M.M.; Funding acquisition, J.M.M

## REFERENCES

1. Peake, J.D., and Noguchi, E. (2022). Fanconi anemia: current insights regarding epidemiology, cancer, and DNA repair. Hum Genet 141, 1811–1836. 10.1007/s00439-022-02462-9.

2. Semlow, D.R., and Walter, J.C. (2021). Mechanisms of Vertebrate DNA Interstrand Cross-Link Repair. Annu Rev Biochem 90, 107–135. 10.1146/annurev-biochem-080320-112510.

3. Garcia-Higuera, I., Taniguchi, T., Ganesan, S., Meyn, M.S., Timmers, C., Hejna, J., Grompe, M., and D’Andrea, A.D. (2001). Interaction of the Fanconi anemia proteins and BRCA1 in a common pathway. Mol Cell 7, 249–262. 10.1016/s1097-2765(01)00173-3.

4. Smogorzewska, A., Matsuoka, S., Vinciguerra, P., McDonald, E.R., Hurov, K.E., Luo, J., Ballif, B.A., Gygi, S.P., Hofmann, K., D’Andrea, A.D., et al. (2007). Identification of the FANCI protein, a monoubiquitinated FANCD2 paralog required for DNA repair. Cell 129, 289–301. 10.1016/j.cell.2007.03.009.

5. Hira, A., Yoshida, K., Sato, K., Okuno, Y., Shiraishi, Y., Chiba, K., Tanaka, H., Miyano, S., Shimamoto, A., Tahara, H., et al. (2015). Mutations in the gene encoding the E2 conjugating enzyme UBE2T cause Fanconi anemia. Am J Hum Genet 96, 1001–1007. 10.1016/j.ajhg.2015.04.022.

6. Rickman, K.A., Lach, F.P., Abhyankar, A., Donovan, F.X., Sanborn, E.M., Kennedy, J.A., Sougnez, C., Gabriel, S.B., Elemento, O., Chandrasekharappa, S.C., et al. (2015). Deficiency of UBE2T, the E2 Ubiquitin Ligase Necessary for FANCD2 and FANCI Ubiquitination, Causes FA-T Subtype of Fanconi Anemia. Cell Rep 12, 35–41. 10.1016/j.celrep.2015.06.014.

7. Knipscheer, P., Räschle, M., Smogorzewska, A., Enoiu, M., Ho, T.V., Schärer, O.D., Elledge, S.J., and Walter, J.C. (2009). The Fanconi Anemia Pathway Promotes Replication-Dependent DNA Interstrand Cross-Link Repair. Science 326, 1698–1701. 10.1126/science.1182372.

8. Klein Douwel, D., Boonen, R.A.C.M., Long, D.T., Szypowska, A.A., Räschle, M., Walter, J.C., and Knipscheer, P. (2014). XPF-ERCC1 acts in Unhooking DNA interstrand crosslinks in cooperation with FANCD2 and FANCP/SLX4. Mol Cell 54, 460–471. 10.1016/j.molcel.2014.03.015.

9. Wang, A.T., Sengerová, B., Cattell, E., Inagawa, T., Hartley, J.M., Kiakos, K., Burgess-Brown, N.A., Swift, L.P., Enzlin, J.H., Schofield, C.J., et al. (2011). Human SNM1A and XPF-ERCC1 collaborate to initiate DNA interstrand cross-link repair. Genes Dev 25, 1859–1870. 10.1101/gad.15699211.

10. Niedernhofer, L.J., Odijk, H., Budzowska, M., van Drunen, E., Maas, A., Theil, A.F., de Wit, J., Jaspers, N.G.J., Beverloo, H.B., Hoeijmakers, J.H.J., et al. (2004). The structure-specific endonuclease Ercc1-Xpf is required to resolve DNA interstrand cross-link-induced double-strand breaks. Mol Cell Biol 24, 5776–5787. 10.1128/MCB.24.13.5776-5787.2004.

11. Bogliolo, M., Schuster, B., Stoepker, C., Derkunt, B., Su, Y., Raams, A., Trujillo, J.P., Minguillón, J., Ramírez, M.J., Pujol, R., et al. (2013). Mutations in ERCC4, encoding the DNA-repair endonuclease XPF, cause Fanconi anemia. Am J Hum Genet 92, 800–806. 10.1016/j.ajhg.2013.04.002.

12. Schlacher, K., Wu, H., and Jasin, M. (2012). A Distinct Replication Fork Protection Pathway Connects Fanconi Anemia Tumor Suppressors to RAD51-BRCA1/2. Cancer Cell 22, 106–116. 10.1016/j.ccr.2012.05.015.

13. Schlacher, K., Christ, N., Siaud, N., Egashira, A., Wu, H., and Jasin, M. (2011). Double-Strand Break Repair-Independent Role for BRCA2 in Blocking Stalled Replication Fork Degradation by MRE11. Cell 145, 529–542. 10.1016/j.cell.2011.03.041.

14. Rogers, C.B., Kram, R.E., Lin, K., Myers, C.L., Sobeck, A., Hendrickson, E.A., and Bielinsky, A.-K. (2023). Fanconi anemia-associated chromosomal radial formation is dependent on POLθ-mediated alternative end joining. Cell Reports 42, 112428. 10.1016/j.celrep.2023.112428.

15. Zellweger, R., Dalcher, D., Mutreja, K., Berti, M., Schmid, J.A., Herrador, R., Vindigni, A., and Lopes, M. (2015). Rad51-mediated replication fork reversal is a global response to genotoxic treatments in human cells. Journal of Cell Biology 208, 563–579. 10.1083/jcb.201406099.

16. Liu, W., Krishnamoorthy, A., Zhao, R., and Cortez, D. (2020). Two replication fork remodeling pathways generate nuclease substrates for distinct fork protection factors. Sci. Adv. 6, eabc3598. 10.1126/sciadv.abc3598.

17. Thangavel, S., Berti, M., Levikova, M., Pinto, C., Gomathinayagam, S., Vujanovic, M., Zellweger, R., Moore, H., Lee, E.H., Hendrickson, E.A., et al. (2015). DNA2 drives processing and restart of reversed replication forks in human cells. Journal of Cell Biology 208, 545–562. 10.1083/jcb.201406100.

18. Lemaçon, D., Jackson, J., Quinet, A., Brickner, J.R., Li, S., Yazinski, S., You, Z., Ira, G., Zou, L., Mosammaparast, N., et al. (2017). MRE11 and EXO1 nucleases degrade reversed forks and elicit MUS81-dependent fork rescue in BRCA2-deficient cells. Nat Commun 8, 860. 10.1038/s41467-017-01180-5.

19. Ceballos, S.J., and Heyer, W.-D. (2011). Functions of the Snf2/Swi2 family Rad54 motor protein in homologous recombination. Biochim Biophys Acta 1809, 509–523. 10.1016/j.bbagrm.2011.06.006.

20. Petukhova, G., Stratton, S., and Sung, P. (1998). Catalysis of homologous DNA pairing by yeast Rad51 and Rad54 proteins. Nature 393, 91–94. 10.1038/30037.

21. Essers, J., Hendriks, R.W., Swagemakers, S.M., Troelstra, C., de Wit, J., Bootsma, D., Hoeijmakers, J.H., and Kanaar, R. (1997). Disruption of mouse RAD54 reduces ionizing radiation resistance and homologous recombination. Cell 89, 195–204. 10.1016/s0092-8674(00)80199-3.

22. Golub, E.I., Kovalenko, O.V., Gupta, R.C., Ward, D.C., and Radding, C.M. (1997). Interaction of human recombination proteins Rad51 and Rad54. Nucleic Acids Res 25, 4106–4110. 10.1093/nar/25.20.4106.

23. Swagemakers, S.M., Essers, J., de Wit, J., Hoeijmakers, J.H., and Kanaar, R. (1998). The human RAD54 recombinational DNA repair protein is a double-stranded DNA-dependent ATPase. J Biol Chem 273, 28292–28297. 10.1074/jbc.273.43.28292.

24. Mason, J.M., Dusad, K., Wright, W.D., Grubb, J., Budke, B., Heyer, W.-D., Connell, P.P., Weichselbaum, R.R., and Bishop, D.K. (2015). RAD54 family translocases counter genotoxic effects of RAD51 in human tumor cells. Nucleic Acids Research 43, 3180–3196. 10.1093/nar/gkv175.

25. Sigurdsson, S., Van Komen, S., Petukhova, G., and Sung, P. (2002). Homologous DNA pairing by human recombination factors Rad51 and Rad54. J Biol Chem 277, 42790–42794. 10.1074/jbc.M208004200.

26. Crickard, J.B., Moevus, C.J., Kwon, Y., Sung, P., and Greene, E.C. (2020). Rad54 Drives ATP Hydrolysis-Dependent DNA Sequence Alignment during Homologous Recombination. Cell 181, 1380–1394.e18. 10.1016/j.cell.2020.04.056.

27. Van Komen, S., Petukhova, G., Sigurdsson, S., Stratton, S., and Sung, P. (2000). Superhelicity-driven homologous DNA pairing by yeast recombination factors Rad51 and Rad54. Mol Cell 6, 563–572. 10.1016/s1097-2765(00)00055-1.

28. Ristic, D., Wyman, C., Paulusma, C., and Kanaar, R. (2001). The architecture of the human Rad54-DNA complex provides evidence for protein translocation along DNA. Proc Natl Acad Sci U S A 98, 8454–8460. 10.1073/pnas.151056798.

29. Alexiadis, V., and Kadonaga, J.T. (2002). Strand pairing by Rad54 and Rad51 is enhanced by chromatin. Genes Dev 16, 2767–2771. 10.1101/gad.1032102.

30. Jaskelioff, M., Van Komen, S., Krebs, J.E., Sung, P., and Peterson, C.L. (2003). Rad54p is a chromatin remodeling enzyme required for heteroduplex DNA joint formation with chromatin. J Biol Chem 278, 9212–9218. 10.1074/jbc.M211545200.

31. Li, X., and Heyer, W.-D. (2009). RAD54 controls access to the invading 3’-OH end after RAD51-mediated DNA strand invasion in homologous recombination in Saccharomyces cerevisiae. Nucleic Acids Res 37, 638–646. 10.1093/nar/gkn980.

32. Wesoly, J., Agarwal, S., Sigurdsson, S., Bussen, W., Van Komen, S., Qin, J., van Steeg, H., van Benthem, J., Wassenaar, E., Baarends, W.M., et al. (2006). Differential contributions of mammalian Rad54 paralogs to recombination, DNA damage repair, and meiosis. Mol Cell Biol 26, 976–989. 10.1128/MCB.26.3.976-989.2006.

33. Selemenakis, P., Sharma, N., Uhrig, M.E., Katz, J., Kwon, Y., Sung, P., and Wiese, C. (2022). RAD51AP1 and RAD54L Can Underpin Two Distinct RAD51-Dependent Routes of DNA Damage Repair via Homologous Recombination. Front Cell Dev Biol 10, 866601. 10.3389/fcell.2022.866601.

34. Uhrig, M.E., Sharma, N., Maxwell, P., Gomez, J., Selemenakis, P., Mazin, A.V., and Wiese, C. (2024). Disparate requirements for RAD54L in replication fork reversal. Nucleic Acids Res 52, 12390–12404. 10.1093/nar/gkae828.

35. Mali, P., Yang, L., Esvelt, K.M., Aach, J., Guell, M., DiCarlo, J.E., Norville, J.E., and Church, G.M. (2013). RNA-guided human genome engineering via Cas9. Science 339, 823–826. 10.1126/science.1232033.

36. Mason, J.M., Chan, Y.-L., Weichselbaum, R.W., and Bishop, D.K. (2019). Non-enzymatic roles of human RAD51 at stalled replication forks. Nat Commun 10, 4410. 10.1038/s41467-019-12297-0.

37. Roy, I.M., Nadar, P.S., and Khurana, S. (2021). Neutral Comet Assay to Detect and Quantitate DNA Double-Strand Breaksin Hematopoietic Stem Cells. Bio Protoc 11, e4130. 10.21769/BioProtoc.4130.

38. Gyori, B.M., Venkatachalam, G., Thiagarajan, P.S., Hsu, D., and Clement, M.-V. (2014). OpenComet: an automated tool for comet assay image analysis. Redox Biol 2, 457–465. 10.1016/j.redox.2013.12.020.

39. Newell, A.E.H., Akkari, Y.M.N., Torimaru, Y., Rosenthal, A., Reifsteck, C.A., Cox, B., Grompe, M., and Olson, S.B. (2004). Interstrand crosslink-induced radials form between non-homologous chromosomes, but are absent in sex chromosomes. DNA Repair (Amst) 3, 535–542. 10.1016/j.dnarep.2004.01.011.

40. Schubert, L., Hendriks, I.A., Hertz, E.P.T., Wu, W., Sellés-Baiget, S., Hoffmann, S., Viswalingam, K.S., Gallina, I., Pentakota, S., Benedict, B., et al. (2022). SCAI promotes error-free repair of DNA interstrand crosslinks via the Fanconi anemia pathway. EMBO Rep 23, e53639. 10.15252/embr.202153639.

41. Amunugama, R., Willcox, S., Wu, R.A., Abdullah, U.B., El-Sagheer, A.H., Brown, T., McHugh, P.J., Griffith, J.D., and Walter, J.C. (2018). Replication Fork Reversal during DNA Interstrand Crosslink Repair Requires CMG Unloading. Cell Rep 23, 3419–3428. 10.1016/j.celrep.2018.05.061.

42. Mutreja, K., Krietsch, J., Hess, J., Ursich, S., Berti, M., Roessler, F.K., Zellweger, R., Patra, M., Gasser, G., and Lopes, M. (2018). ATR-Mediated Global Fork Slowing and Reversal Assist Fork Traverse and Prevent Chromosomal Breakage at DNA Interstrand Cross-Links. Cell Rep 24, 2629–2642.e5. 10.1016/j.celrep.2018.08.019.

43. Hanada, K., Budzowska, M., Davies, S.L., van Drunen, E., Onizawa, H., Beverloo, H.B., Maas, A., Essers, J., Hickson, I.D., and Kanaar, R. (2007). The structure-specific endonuclease Mus81 contributes to replication restart by generating double-strand DNA breaks. Nat Struct Mol Biol 14, 1096–1104. 10.1038/nsmb1313.

