## Supplemental Figures for "RAD54L promotes nascent DNA degradation and radial chromosome formation in FANC-deficient cells"

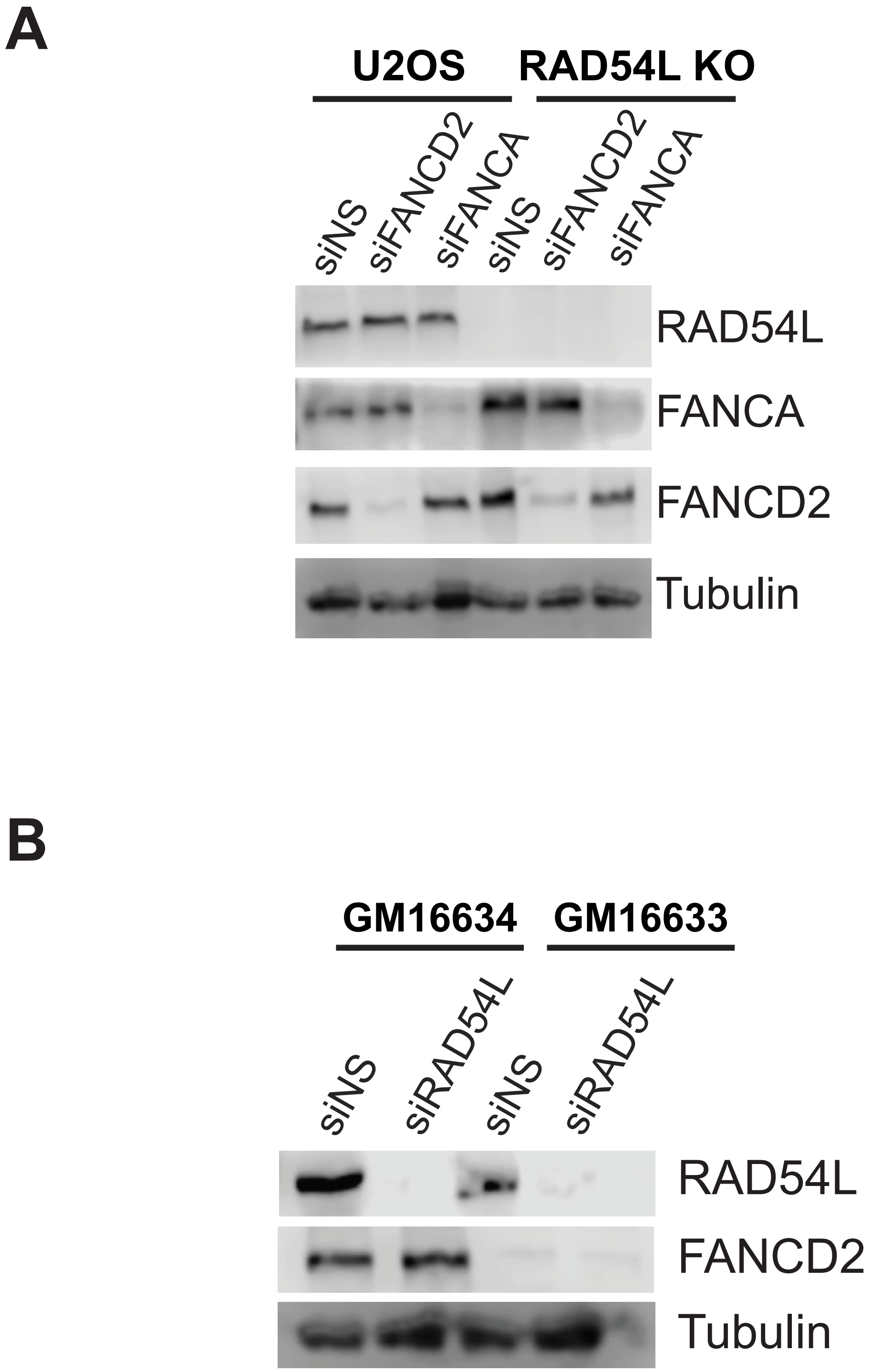


**Supplemental Figure 1**. **Westerns depicting protein levels in indicated cell lines**. (A) Western blot in U2OS and RAD54L knockout cell lines showing siRNA knockdown of FANCD2 and FANCA. Tubulin as loading control. (B) Western blot showing RAD54L siRNA knockdown and FANCD2 knockout in GM16633 and GM16634. Tubulin used as a loading control.

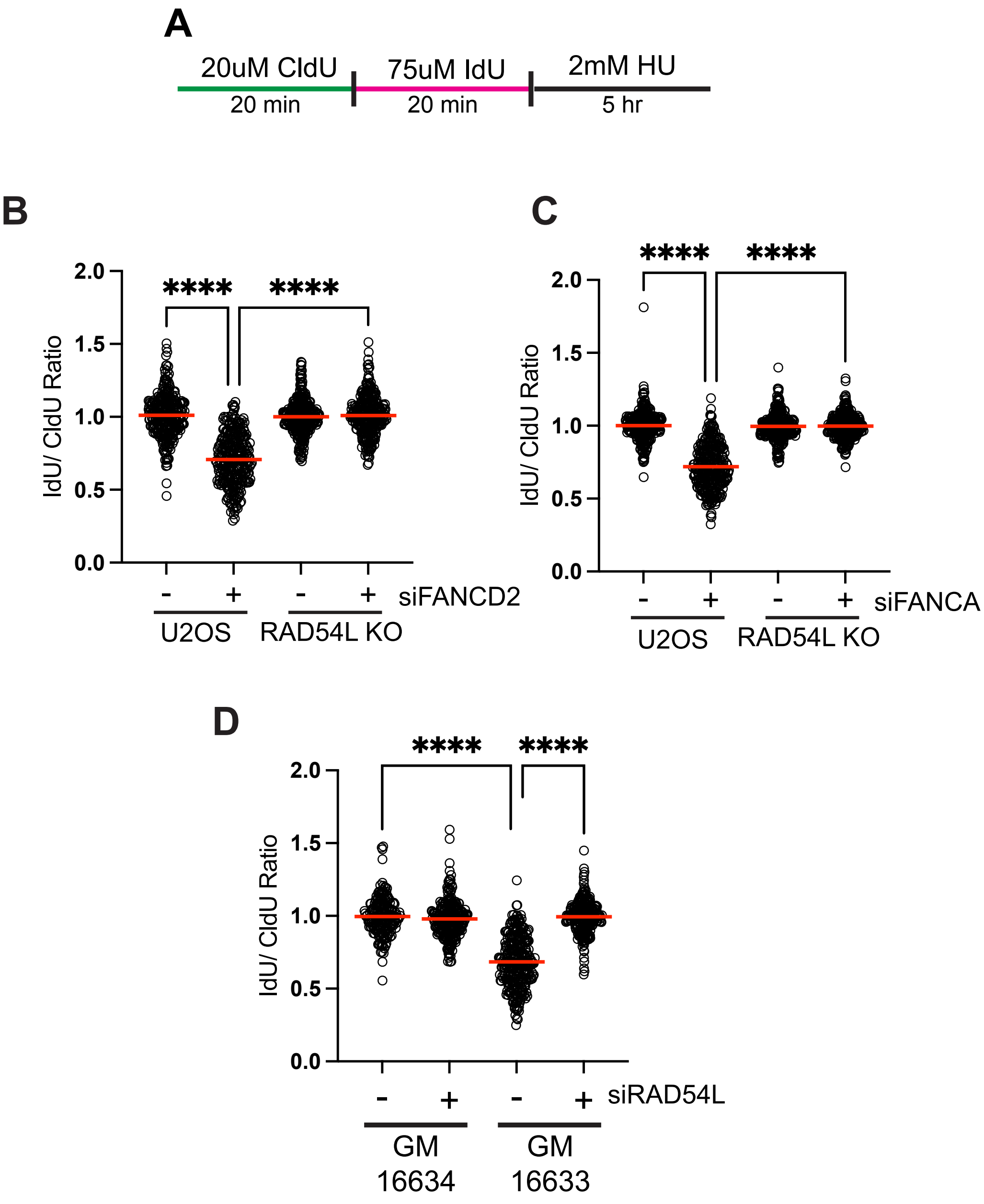


**Supplemental Figure 2. Depletion of RAD54L rescues fork degradation in FANCD2 and FANCA deficient cells following treatment with hydroxyurea (A)** Schematic representing replication fiber assay **(B)** Dot blot showing IdU/CldU ratios of replication fibers. Red line represents the mean from two independent experiments *****p*<0.0001, ANOVA, Tukey HSD. (**C)** Dot blot showing IdU/CldU ratios of replication fibers. Red line represents the mean from two independent experiments. *****p*<0.0001, ANOVA, Tukey HSD. (**D)** Dot blot showing IdU/CldU ratios of replication fibers. Red line represents the mean from two independent experiments. *****p*<0.0001, ANOVA, Tukey HSD.
